# An extensive computational approach to inhibit MSP-1 of *P.vivax* elucidates further horizon in the establishment next generation therapeutics against malaria

**DOI:** 10.1101/2020.05.02.073973

**Authors:** Md Ohedul Islam, Parag Palit, Jakaria Shawon, Md Kamrul Hasan, Mustafa Mahfuz, Tahmeed Ahmed, Dinesh Mondal

**Affiliations:** International Centre for Diarrhoeal Disease Research, Bangladesh; Department of Biochemistry and Molecular Biology, Tejgaon College, Bangladesh

**Keywords:** merozoite surface protein-1, computational approach, siRNA designing, molecular docking

## Abstract

Malaria represents a life-threatening disease caused by the obligate intra-erythrocytic protozoa of the *Plasmodium* genus, exerting a sinister global health burden and accounting for approximately 660,000 deaths annually. Additionally, 219 million new cases are reported each year, most of which result from the growing issue of artemisinin resistance shown by the *Plasmodium* parasite. Much of the research done for the purpose of development of therapeutics against malaria has traditionally been focused on *Plasmodium falciparum*, which is responsible for majority of the cases of mortality due to malaria, *Plasmodium vivax* is also known to contribute greatly towards the malaria relate morbidities particularly in vivax endemic areas. In this study, we have used two different computational approaches aimed at establishing newer concepts towards the development of advanced therapeutics against vivax malaria by targeting the surface antigen, merozoite surface protein-1 (MSP-1). In-silico approach involving computational siRNA designing against MSP-1 resulted in a total of four candidate siRNAs being rationally validated following corroboration with a plethora of algorithms. Additionally, molecular docking analysis unraveled a total of three anti-parasitic peptides. These peptides namely: AP02283, AP02285 and AP00101 were found to exhibit considerable binding affinity with MSP-1 of *P.vivax*, thus providing an apparent indication of their anti-malarial property and affirming their potency to be used as novel molecules for development of next generation anti-malarials. However, irrespective of the prospective magnitude of these in-silico findings, the results require extensive validation by further rigorous laboratory experiments involving both in-vitro and in-vivo approaches.

## Introduction

Malaria, a serious global health burden with an estimated number of 219 million new incidences of morbidity occurring each year and resulting 660,000 deaths annually [1]. It represents a life-threatening disease caused by the obligate intra-erythrocytic protozoa of the *Plasmodium* genus of parasites that are transmitted to people primarily through the bites of infected female *Anopheles* mosquitoes, with high transmission rates being prevalent in the regions of sub-Saharan Africa, Papua New Guinea, and the South Pacific islands [2]. Infections can also either be congenitally acquired or may occur through exposure to infected blood products, a phenomenon known as transfusion malaria [3]. Humans can be infected by one or more of the five species of the *Plasmodium* genus, including: *P. falciparum, P. vivax, P. ovale, P. malariae, and P. knowlesi*, among which *P.vivax* is the most prevalent in different parts of Asia and the Latin American subcontinent and there are a few places where *P. vivax* is transmitted exclusively [4].Unlike *P. falciparum, P. vivax* infections are marked by low blood-stage parasitemia with gametocytes emerging before the manifestation of illness and dormant liver stages, causing relapses [6].

The development of an effective anti-malarial therapy has become a major public health challenge. Acute clinical malaria, which is often life-threatening is associated with replication of the asexual blood stage parasite in circulating erythrocytes [7].. Advances towards the development of new anti-malarials have revealed the *Plasmodium* surface antigen called merozoite surface protein–1 (MSP-1) to be considered as a potent target for development of new anti-malarial therapeutics [9]-[10].

MSP-1 is an antigen which is abundantly expressed on surface of mature merozoites [11] and has been shown to participate in the parasite invasion of the erythrocyte [12]. Structural studies on the gene indicate that parts of the molecule are well-conserved (Holder et al., 1992). Strong evidences suggest that MSP1-specific immune response can protect against the blood-stage challenge [13] and certain mAbs to MSP-I in a number of different *Plasmodium* species have been shown to inhibit the growth of the parasite in vitro [14]-[15] or protect on passive transfer in vivo [16]-[17].There is an urgent requirement for effective therapeutics against malaria caused by the *Plasmodium* parasite, *P.vivax*, and proteins on the surface of the malarial merozoite are potent targets for development of anti-malarial therapeutic owing to their capability overt to antibody generated by the host defense system [18].

A newly emerging therapeutic approach a number of diseases involves utilizing the means of post transcriptional gene silencing through the use of siRNA molecules [23]. The central role of siRNA is post transcriptional gene silencing by interfering with the protein and their basic mechanism is to bind and promote the degradation of messenger RNA (mRNA) at specific sequences [24].. The use of nanoparticles like polyethyleneimine or small cationic peptides can facilitate the introduction of the siRNA into the cell for specific knockdown of a gene of interest [26].

On the contrary, the use of anti-microbial peptides represents a novel post-therapeutic strategy to combat microbial infestations and reduce the incidences of the development of anti-microbial resistance. Antimicrobial peptides (AMPs) are host defense molecules universal in the innate immune systems of both invertebrates and vertebrates [27]-[28]. They form the first line of host defense against pathogenic infections and are key components of the innate immune system. Because these ancient molecules remain potent after millions of years, they are regarded as important templates for developing a new generation of antimicrobials to combat numerous antibiotic resistant superbugs and parasites [29]-[30]. Host insects and general arthropods, which act as reservoirs for such parasites have developed a way to coexist with such parasites. These organisms are thus a promising source for the prospection of anti-parasitic compounds, as alternative methods for the treatment of protozoa-related diseases [31]. Among the molecules already isolated and investigated, there are proteins and peptides with high activity against parasites, able to inhibit parasite activity in different stages of development [32],[33],[34]. Although, studies are still taking their first steps, initial results show new perspectives on the treatment of parasitic diseases.

In this current study, we have designed several therapeutic approaches targeting MSP-1 of *P.vivax*, with the intention to inhibit MSP-1 and establish new approaches for anti-malarial therapy against vivax malaria. Among these next generation therapeutics, such as gene silencing by introduction of functional siRNA and administration of purified antiparasitic peptides can serve as convenient alternatives for development of post therapeutic options for vivax malaria.

## Methods and Materials

An outline of the whole methodology undertaken for this study has been portrayed in Figure 1

**Figure 01:**
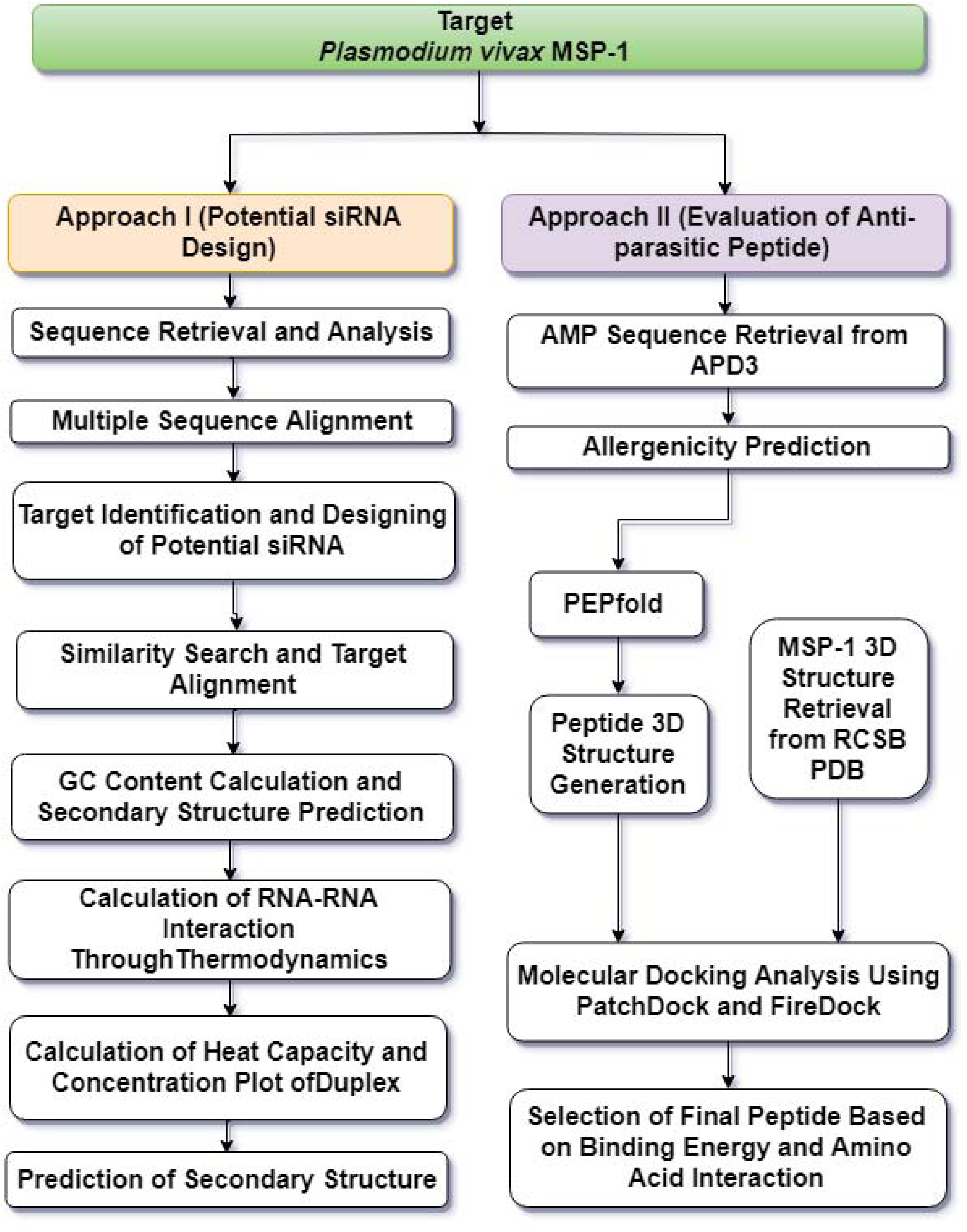
Flowchart summarizing the protocols undertaken to complete this molecular evaluation study

### Approach I: Designing of potential siRNA candidate

#### Sequence Retrieval and Analysis and Multiple Sequence Alignment

Complete cds of msp-1 gene sequences from a total of 92 isolates of *P.vivax* were retrieved from the gene bank database (Accessed on March 2018) available at National Centre for Biotechnological Information (https://www.ncbi.nlm.nih.gov/). Subsequent multiple sequence alignment was done using ClastalO (https://www.ebi.ac.uk/Tools/msa/clustalo/).

#### Target identification and designing of potential siRNA

siDirect 2.0 (http://sidirect2.rnai.jp/) an efficient and target-specific siRNA designing tool was used for target identification and siRNA designing [43]. In siDirect 2.0 Ui-Tei, Amarzguioui, Reynolds rules and melting temperature below 21.5 °C were considered as absolute parameters for potential siRNA duplex formation. Additionally, other components were also taken into account on the concept of algorithms, as illustrated in **Table 01**.

**Table 01:** Algorithms and rules for rational design of siRNA molecules

| Ui–Tei rules | Amarzguioui rules | Reynolds rules |
| --- | --- | --- |
| A/U at the 5' terminus of the sense strand | Duplex end A/U differential >0 strong binding of 5' sense strand | Each rule is assigned a score which is summed up to a total duplex score to improve the efficiency of siRNA |
| G/C at the 5' terminus of the antisense strand | No U at position 1. Presence of A at position 6 |  |
| At least 4 A/U residues in the 5' terminal 7 bp of sense strand | Weak binding of 3' sense strand. No G at position 19 |  |
| No GC stretch longer than 9 nucleotides. |  |  |

#### Similarity Search and Target Alignment

BLAST tool (http://www.ncbi.nlm.nih.gov/blast) was used to check any off-target sequence resemblance in the genome of other non-targeted organism, especially that of humans. BLAST was used against whole Genebank database by applying expected thresholds value 10 and BLOSUM 62 matrix as parameter.

#### GC Content Calculation and Secondary Structure Prediction

DNA/RNA GC content calculator (http://www.endmemo.com/bio/gc.php) was used to assess the GC content of the predicted siRNA. Consequently, the secondary structure and free energy of folding of siRNA was predicted using mfold server (http://unafold.rna.albany.edu/?q=mfold) [44].

#### Calculation of RNA-RNA Interaction through Thermodynamics

To investigate the thermodynamics of the interaction between the predicted siRNA and their target, RNAcofold program (http://rna.tbi.univie.ac.at/cgi-bin/RNAWebSuite/RNAcofold) was used [45]. This program computes the hybridization energy and base pairing from two RNA sequences. RNAcofold follows the functional algorithm of McCaskill’s partition to compute probabilities of base pairing, realistic communication energies and equilibrium concentrations of duplex structures.

#### Calculation of Heat Capacity and Concentration Plot of Duplex

DINAMelt server has been used to calculate the heat capacity plot and concentration plot (http://unafold.rna.albany.edu//?q=DINAMelt/Hybrid2) for the predicted siRNA [46]. The heat capacity (Cp) is plotted as a function of temperature, with the melting temperature T_m_ (Cp) indicated. The detailed heat capacity plot also shows the contributions of each species to the ensemble heat capacity, whereas concentration plot—T_m_(Conc.), the point at which the concentration of double-stranded molecules of one-half of its maximum value defines the melting temperature T_m_(Conc.).

#### Prediction of Secondary Structure

The secondary structure formed between the target sequence and siRNA was predicted using RNAstructure (https://rna.urmc.rochester.edu/RNAstructureWeb/Servers/Predict1/Predict1.html) web server [47]. This tool generates secondary structures of siRNA and target interaction by calculating a partition function, predicting a maximum free energy structure along with maximum expected accuracy.

### Approach II: Evaluation of Anti-Parasitic Peptides as Anti-malarial

#### AMP sequence retrieval and filtering

APD3 database (http://aps.unmc.edu/AP/main.php), a comprehensive database for all the anti-parasitic peptides, including and anti-malarial peptides [28]. Amino acid sequences for a total of 78 anti-parasitic peptides and 25 anti-malarial peptides (accessed on March 2018) had been retrieved. The antimalarial peptides that were found to be common or overlap with the anti-parasitic peptides were discarded from the study.

#### Allergenicity prediction

It is very important for any foreign peptide sequence to be non-allergenic considering the human immune system and its scrupulous defense mechanism [48]. Peptides that could elicit possible allergic response were thus avoided by filtering them on the basis of their allergenicity. For this purpose, the peptide sequences were subjected to Algpred (http://crdd.osdd.net/raghava/algpred/index.html), an online based tool that allows prediction of allergens based on similarity of known epitope with any region of peptide sequence.[49] A single parameter was set to Hybrid option that allows the server to predict allergen using combined approach (SVMc + IgE epitope + ARPs BLAST + MAST).

#### Molecular Docking of Anti-parasitic peptide and MSP-1

##### Generation of 3D structures of the peptides

Non-allergic peptide sequences were selected as input in Pepfold server (http://bioserv.rpbs.univ-paris-diderot.fr/services/PEP-FOLD3/) in order to generate peptide 3d structures. In Pepfold server the number of simulations used was 200 and sOPEP sort model was set as a default parameter.

##### MSP-1 3D structure retrieval

Protein structure of MSP-1 (2NPR) was downloaded from the Protein Data Bank (https://www.rcsb.org/) database. This structure (2NPR) is basically a 15 kDa subunit MSP-1 (19) cleaved from the C-terminal MSP-1 sub-unit MSP-1(42), during red blood cell invasion. 2NPR includes 20 states of the MSP-1 (19) and was saved in the pdb file format by using PyMOL state 1 for downstream molecular docking analysis

##### Molecular Docking Simulation

Molecular docking of MSP-1 (2NPR) against selected peptides was done using PatchDock (https://bioinfo3d.cs.tau.ac.il/PatchDock/) server along with FireDock docking server (http://bioinfo3d.cs.tau.ac.il/FireDock/), a refinement and rescoring tool used to refine the PatchDock output. MSP-1(19) structure (2NPR) along with single peptide structure in PDB format were uploaded in PatchDock server for initial calculation and the top 10 resultant docking solutions were directly uploaded to FireDock server for further calculation. Interaction analysis was done after docking to see whether the peptides remain in the binding pocket or not using Accelrys Discovery Studio [50].

## Results

### Therapeutic approach I: siRNA designing against msp-1

#### MSP-1 nucleotide sequence collection and multiple sequence alignment

Complete gene sequences for msp-1 from 92 different isolates of P.vivax were obtained from NCBI Nucleotide. Subsequent multiple sequence analysis using Clustal Omega revealed a total of 17 conserved sequences (shown in supplementary file 2) which were used for further analysis through this approach.

#### Designing siRNA by siDirect

A web based highly effective siRNA computational tool siDirect 2.0 was used in the next step. All 17 conserved sequences were as input sequence to siDirect 2.0 to design efficient, target specific siRNA with decreased probability of off-target silencing. This tool predicted a total of 36 different siRNA molecules against all the 17 conserved sequences. These results were further filtered on the basis of following all three rules governing the sequence preference of siRNA, namely: Ui-Tei, Amarzguioui and Reynolds rules (U, R, A rules). Among all these 36 predicted siRNA molecules, only 13 siRNAs were found to have complied with all the three algorithms completely, i.e-U,R,A rules **(Supplementary table 01)**. Off-target resemblance of siRNAs minimization was confirmed using NCBI BLAST program.

#### GC content of predicted siRNA

GC content of siRNAs is a determinant of the stability of its secondary structure potential and 30-57% of GC content is considered sufficient for the execution of its action [51]. In this study, results from DNA/RNA GC content calculator showed that among 13 predicted siRNA 12 have GC content hold within the range of 33-48% **(Table 04)**.

**Table 02:**
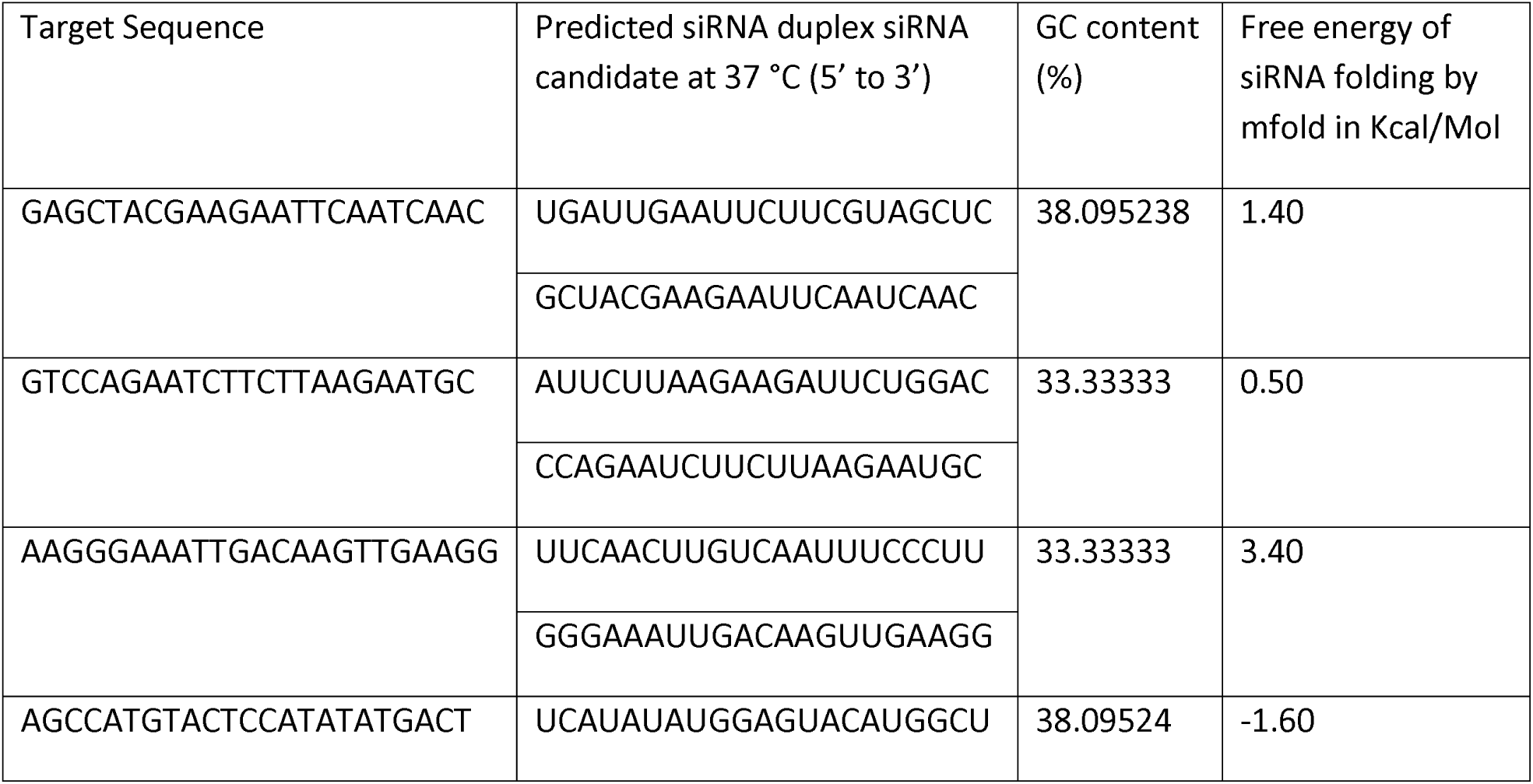
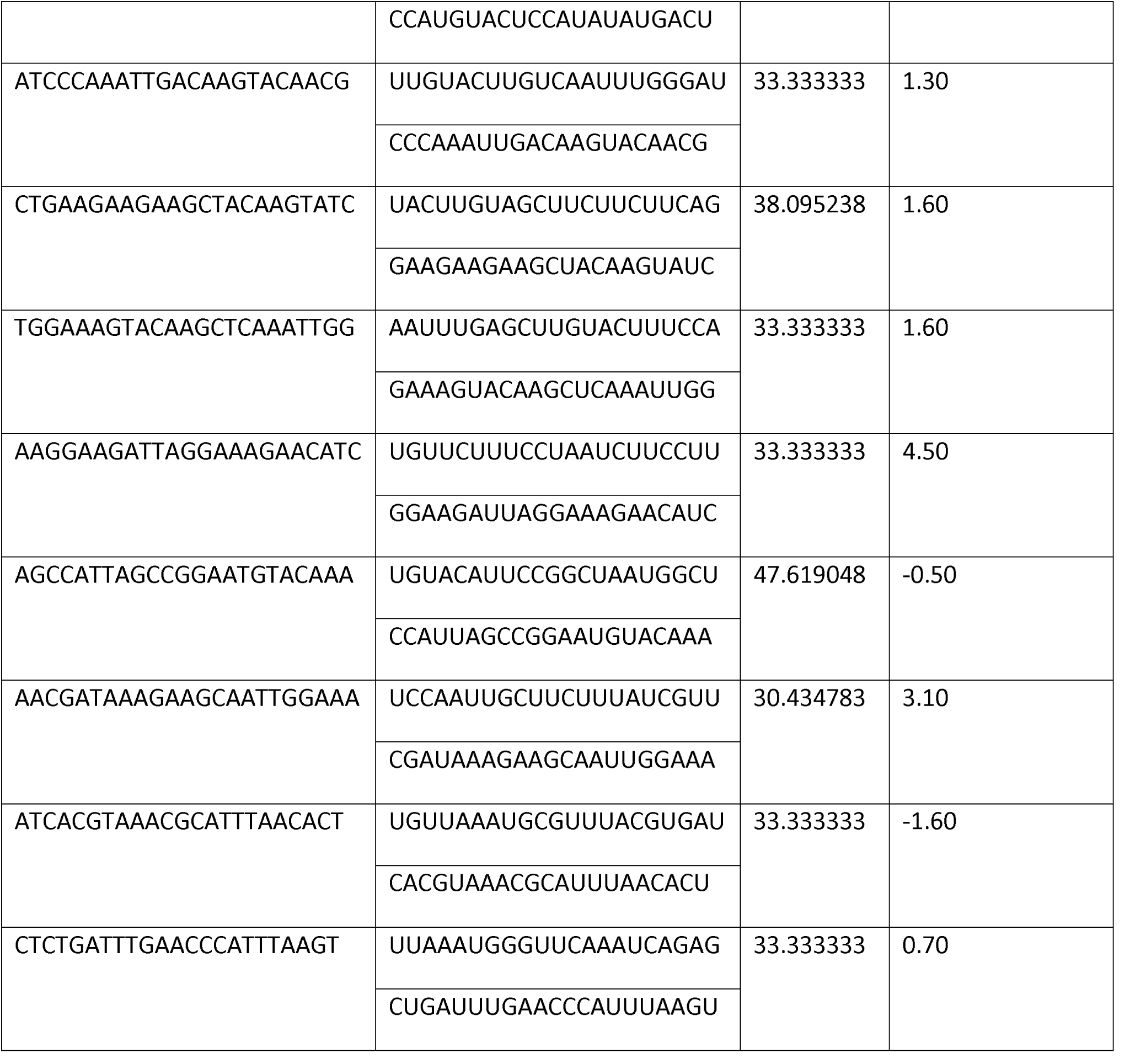
Effective siRNA molecule with GC%, free energy of folding

**Table 03:**
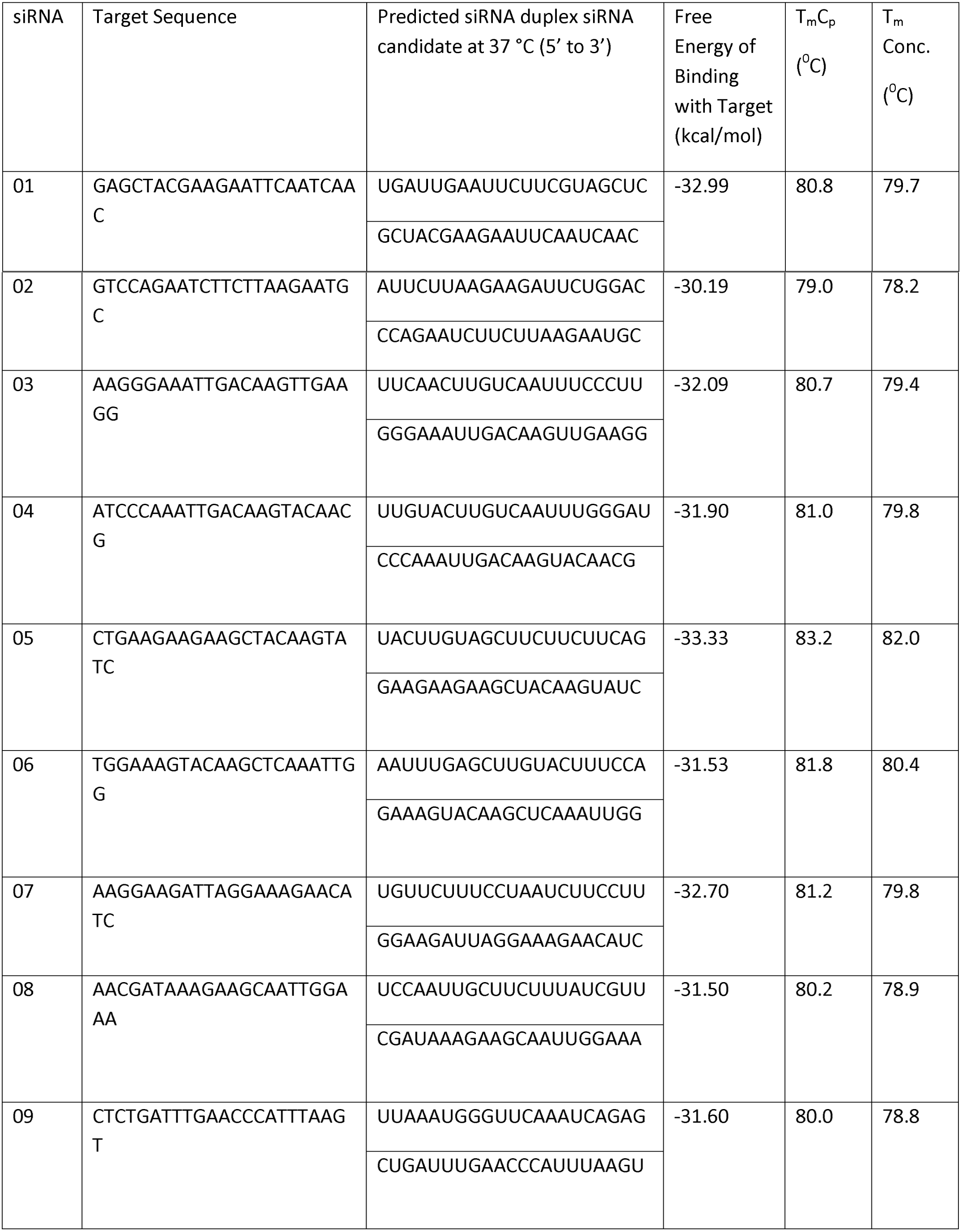
Free energy of binding of siRNA with target, heat stability or heat capability temperature

**Table 04:**
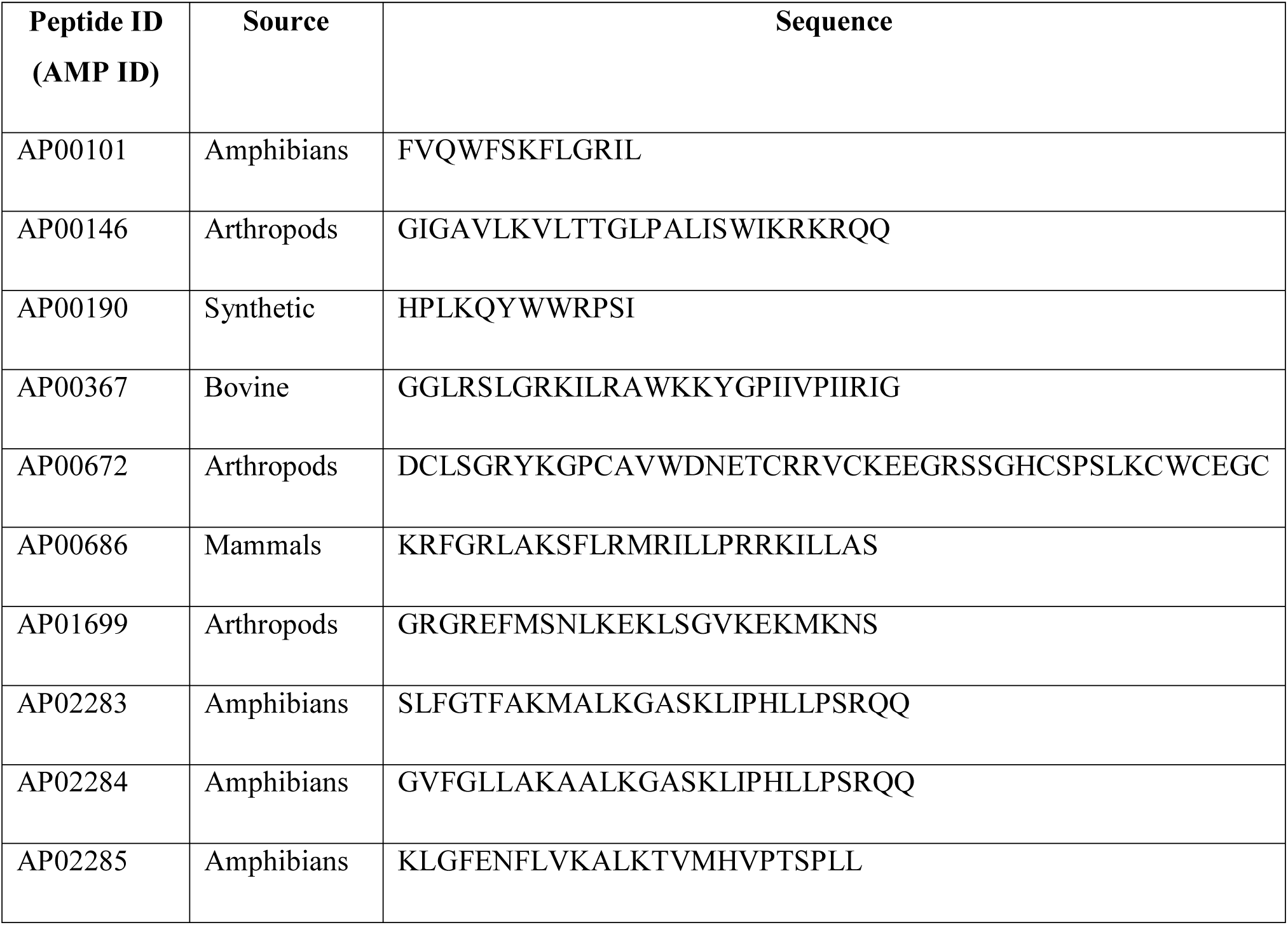
Anti-parasitic peptide sequences retrieved from AMP database having non-allergic characteristics and their sources

#### Secondary structure prediction

Since the function of siRNA is also heavily dependent on molecular structure, considerable efforts are made in computational methods for predicting RNA secondary structure [52]. Minimum free energy (MFE) is considered as benchmark of molecular structural accuracy of the siRNA [53]. Free energy minimization is a long established paradigm in computational structural biology that is based on the assumption that, at equilibrium, the solution to the underlying molecular folding problem is unique, and that the molecule folds into the lowest energy state [54].In this molecular evaluation process, the minimum free energy of folding was calculated **(Table 05)** by using the mfold server to check the stability of the predicted siRNA guide strands [44]. Here, 9 out of 12 predicted siRNA passed this filter with positive free energy. The folded structures are showed in **figure 05.**

**Table 05:**
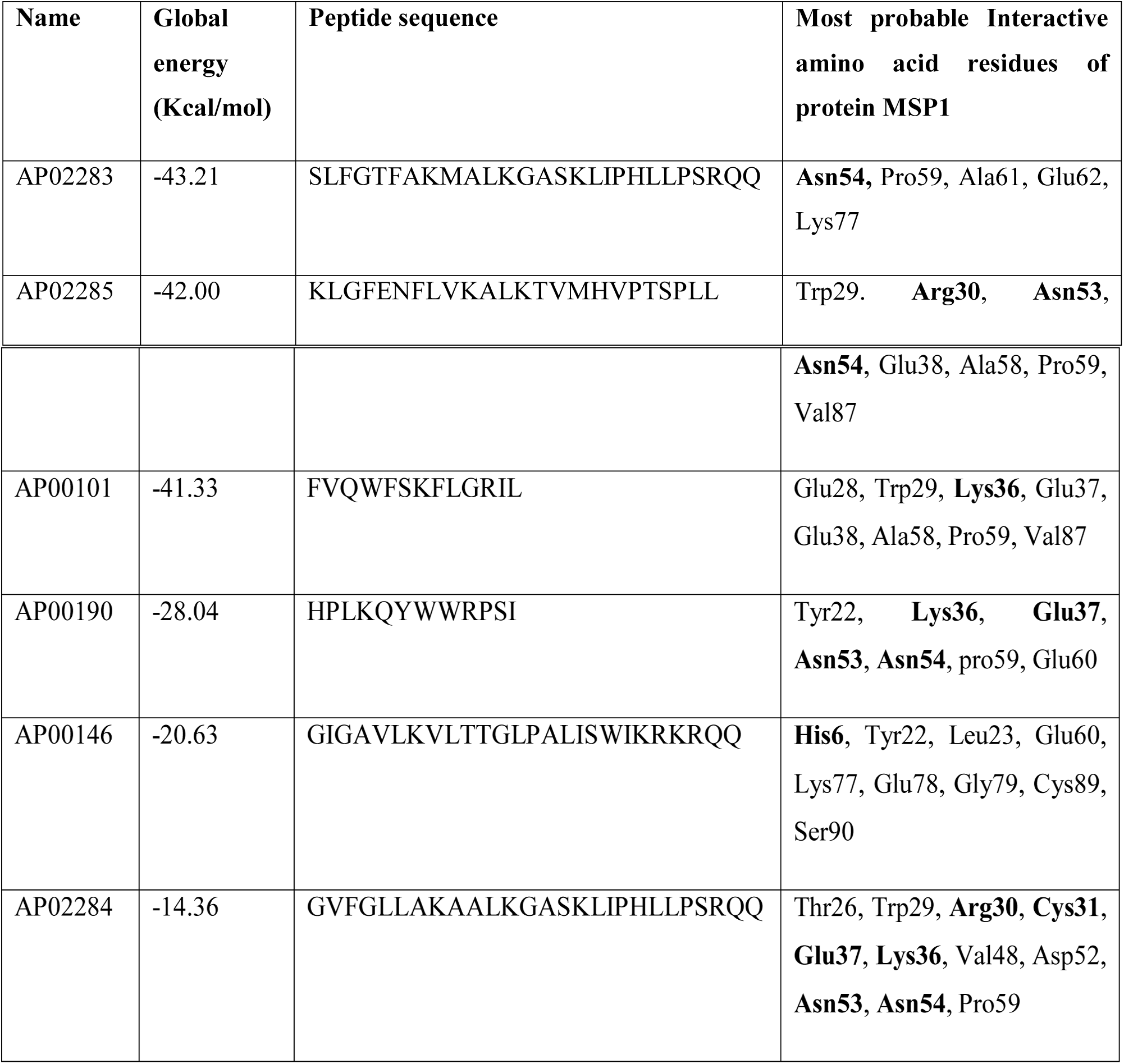
Docking of peptides against 2NPR using PatchDock and Firedock, Global Energy and Interactive Amino Acids. Amino Acids in bold letters are the matching binding amino acids in the msp-1 binding groove.

#### RNA-RNA binding energy calculation

Reliable prediction of RNA–RNA binding energies is crucial for deeper understanding of the binding of the RNAi molecule to the target mRNA. RNAcofold is a program from Vienna RNA webservers, which was used to calculate the hybridization energy and base-pairing pattern of two RNA sequences [55]. The thermodynamics of such RNA–RNA interactions can be understood as the sum of two energy contributions: (1) the energy necessary to ‘open’ the binding site and (2) the energy gained from hybridization [56]. Hence the accurate computational treatment of RNA-RNA binding is integral to the functioning target prediction algorithms. For a single siRNA the program annotates two sequences i.e. sequence A and sequence B and link them with a tag to that specific link. In addition to minimum free energy of heterodimer (ΔGAB), the free energy of the heterodimer (ΔGA and ΔGB) of sequence A and sequence B can be calculated using the following equation [45]- 

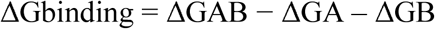

Using this equation free energy of binding (ΔGbinding) is calculated. Such as, for the interaction of target sequence 1 and its predicted siRNA, free energy of binding is- 

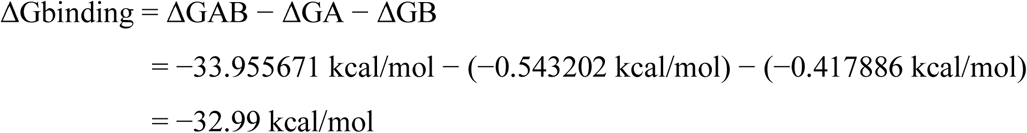

Rest of the target sequences and their predicted siRNA were subjected to RNAcofold and the free binding energy was calculated according to default parameter setting **(Table 05)**. For further use and compatibility, in table 04 all the siRNA was defined as siRNA 01-09.

#### Heat stability Analysis

DINAMelt web server can predict entire equilibrium melting profiles of any as a function of temperature with melting temperature for a hybridized pair of nucleic acids. Heat capacity or Tm (Cp) is expressed as a function of temperature while Tm (Conc) is the temperature at which the temperature of nucleic acid reaches half of its maximum value [46]. It is known that the secondary structure can indeed cause inhibition of RNAi [57]. In this study, all the predicted siRNA showed T_m_ (Cp) and T_m_(Conc.) around 80°C **(Table 05)** which is way above the in vivo temperature (37.4°C).

#### Prediction of secondary structure

Secondary structures of RNAi, including that of siRNAs provide a convenient and computationally tractable estimation not only to structures but also to the thermodynamics of RNA-RNA interaction. RNAstructure program predicts the most stable secondary structures of target-siRNA duplex structure by minimizing the folding free energy. The folded structures are predicted at most convenient temperature that is 37°C. siRNA that bound to its target sequence with highest free energy (kcal/mol) while forming duplex were suggested to be our most potential siRNA. Among all 9 duplexes, siRNA 01, siRNA 05, siRNA 04 and siRNA 07 showed highest free energy of binding that was -31.4, -31.4, -30.0, and -30.0 kcal/mol respectively. These predicted duplex secondary structures are depicted in **figure 06**.

### Therapeutic Approach II: Anti-Parasitic peptide evaluation as Anti-malarial

#### Peptide sequence retrieval and antigenicity prediction

From the APD3 database a total of 101 anti-parasitic and 25 anti-malarial peptides were retrieved. Majority of these peptides were found to be either of amphibian or arthropod origin, which are known to be a major source of antimicrobial peptides [28]. These 25 antimalarial were found to overlap with the list of antimalarial peptides obtained from the database and were subsequently excluded from the study. 76 of the remaining anti-parasitic peptides were tested for allergenicity and among them 10 were found to have non-allergic characteristics and were selected for further analyses **(Table 06)**.

#### Peptide 3D structure generation

PEP-FOLD produces informative report to guide the user through the best model selection [41]. This server predicts a cluster of structures for each query sequence. The cluster ranks are defined according to their sOPEP score and top scored structures were downloaded as pdb for docking analysis.

#### Molecular Docking of Anti-parasitic peptide and MSP-1

PatchDock provided docking complex of peptide-2NPR with refining score. Top 10 complexes were directed to FireDock with refining score for further enhancement and rescoring in terms of global energy (i.e., the binding energy of the solution). In spite of having a substantial global energy, 6 peptide-2NPR complexes were listed as final result and 4 were phased out as they were not found to bind to the intended site of the protein. On the basis of our docking study, all 6 peptides showed good binding energy interacting with at least one exact binding groove amino acids reported earlier by different groups [58]-[59]. We used discovery studio visualizer to find the most probable interactive amino acids of msp-1 **(shown in Table 07)**.

Interactive images of peptide and MSP-1 (19) docking was illustrated via discovery studio visualizer and depicted in **figure 07**. AP02285, AP00190 and AP02284 showed maximum interactions with target amino acid residues as per previous research from other groups related to MSP-1 structure, binding groove and selection of inhibitor.

## Discussion

Among all the species of the *Plasmodium* genus, *P. falciparum* is the most severe and responsible for 90% of all deaths due to malaria[60]. Despite this, *P.vivax* is still responsible for majority cases of morbidity and significant mortality in vivax endemic areas [5]. Regardless of such a large burden of disease, *P. vivax* has always been overlooked and left in the shadow. In this study we aimed at the merozoite surface protein 1 (msp-1) of *Plasmodium vivax* that is primarily responsible for the binding to reticulocyte eventually resulting in RBC invasion [62]. A combined approach involving “computational design of siRNA with RNAi activity against MSP-1 gene” in conjugation with “anti-parasitic peptide-based inhibition of MSP-1 protein” were carried out in attempts to elucidate novel means to restrict the functional activity of MSP-1.

Our study, also comprised of designing siRNA against MSP-1, which represents a promising new approach for producing mRNA specific inhibition [68]-[69]. A candidate siRNA that follows all the rules and algorithms of Ui-Tei, Amarzguioui and Reynolds is supposed to be the most effective[70]. BLAST program was employed to avoid off target activity that can complicate the silencing activity of siRNA and can potentially lead to unwanted toxicities[71]. It is recommended to select siRNA sequences with low GC content, preferably 30% to 57% [51], since there is a considerable reciprocal correlation between GC-content and RNAi activity. Therefore, lower GC content of siRNA exhibits stronger inhibitory effect [72].

The molecular structure of siRNA is crucial for the functional activity and considerable number of computational methods are available for siRNA secondary structure prediction [52]. For RNA molecular structural stability and accuracy, the minimum free energy (MFE) is considered as a standard parameter [53]. The nine predicted siRNA were further considered for analysis and the rest were discarded as negative value of free energy of folding may reduce the accessibility of siRNA specific to its target site.

The thermodynamic outcome of the interaction between the predicted siRNA and the target mRNA can be understood using the sum of the energy necessary to “open” the binding site and the energy gained from hybridization[56]. For the thermodynamic prediction of the RNA-RNA interaction efficiency, all the siRNA was subjected to Vienna RNA webserver. During siRNA designing for a therapeutic purpose, it must be ensured that at a certain temperature the siRNA product is stable and functional in vivo. DINAMelt web server simulates not only the melting of siRNA in solution but also the entire equilibrium melting profiles as a function of temperature. The heat capacity Tm (Cp) works as a function of temperature whereas Tm(Conc.) is the temperature at which the temperature of siRNA reaches half of its maximum value [46]. Regarding heat capacity, our predicted siRNA possesses a strong stability at very high temperatures.

On the other hand, randomly folded siRNA can cause inhibition of its RNAi activity [73] and improper secondary structure of siRNA can lead to RISC complex formation. Moreover, prediction of secondary structure of target-siRNA duplex is important to guide the selection of specific siRNA target site [72]. Secondary structure of target-siRNA duplex gives computationally valuable information for structure design and thermodynamics of RNA-RNA interaction. Finally, 4 predicted siRNA in this study showed reliable duplex secondary structure with a considerable MFE value (Figure 06).

Small peptide can work as a potent inhibitor of many biological process and anti-parasitic peptides (APPs) are not different to this. Among the different sources of APPs, exogenous and synthetic APPs are found to be more potent in inhibitory effect [74]. In this computational experiment, we aimed to check and evaluate the entire library of APPs present in the APD3 database and their activity against MSP-1 as an inhibitor as well as anti-malarial therapeutic agent. The prediction of allergenicity of the peptide sequences downloaded from database is important because allergic diseases are among the most common health issues worldwide [75]. Many allergic reactions are mild, while others can be severe and life threatening. In AlgPreda systematic attempt has been made to integrate various approaches in order to predict allergenic proteins with high accuracy [76]. Our data shows that among the 76 APP assessed, 10 were found to be non-allergen passing all the combined filters of Algpred.

Prior to docking, the MSP-1 (19) 3D structure (2NPR) from PDB database was downloaded and all peptide 3D structure generated by PEP-fold server. PEP-FOLD is based on a Hidden Markov Model derived Structural Alphabet [77] that allows the treatment of both linear and di-sulfide bonded cyclic peptides that range from 9-36 amino acids [41]. In docking, both MSP-1 (19) and peptides pdb structures were used in PatchDock which follows an algorithm inspired by object recognition and image segmentation for molecular dockings. For each peptide-protein docking PatchDock generates a list of potential docked complexes where the candidates are ranked according to a geometric shape complementarity score. Resultant docked complexes along with refining score directed to FireDock to refine and re-score the rigid complexes of peptide-protein docking solutions. The final results were ranked by the global energy value, i.e-the binding energy of the solution). Babon JJ et al. reported that His6, Arg21, Glu27, Lys36, Asn47 and Asn54 of the protein MSP-1 (19) (2NPR) could be the most obvious candidates for interaction with ligands [59]. In our study, we observed that after docking analysis 6 peptides (AP28302, AP02285, AP00101, AP00190, AP00146, and AP02284) interacted with candidate amino acids in MSP-1 with proficient global binding energy (Figure 07). AP02283, AP02285 and AP00101 were found to have the highest global binding energy and interacts specifically with Asn54, Asn54, and Lys36 of MSP-1(19) binding cleft respectively. This observation helped to assume that AP02283, AP02285 and AP00101 binds to the binding cleft of MSP-1(19) and can inhibit reticulocyte binding and efficiently work as MSP-1 inhibitors.

## Conclusion

MSP-1 is a major surface antigen of *Plasmodium* sp. that is known to exhibit substantial antigenicity and is responsible for invasion of erythrocytes and subsequent pathogenicity. These properties open up new horizons for MSP-1 to be used for development of newer anti-malarial therapeutics. However, the findings in this study are nevertheless accompanied with several pitfalls. Possibilities are there of the predicted siRNA exhibiting off-target silencing due to abundant polymorphisms among different ethnicities. Moreover, it is not expected that biotechnological peptides will take over chemical molecules on the development of new drugs. Henceforth, further lab studies on a considerable scale are needed for the validation of these findings. Nevertheless, the outcomes of this study will serve as robust background data for the purpose of such application-based studies in the long run.

**Figure 02:**
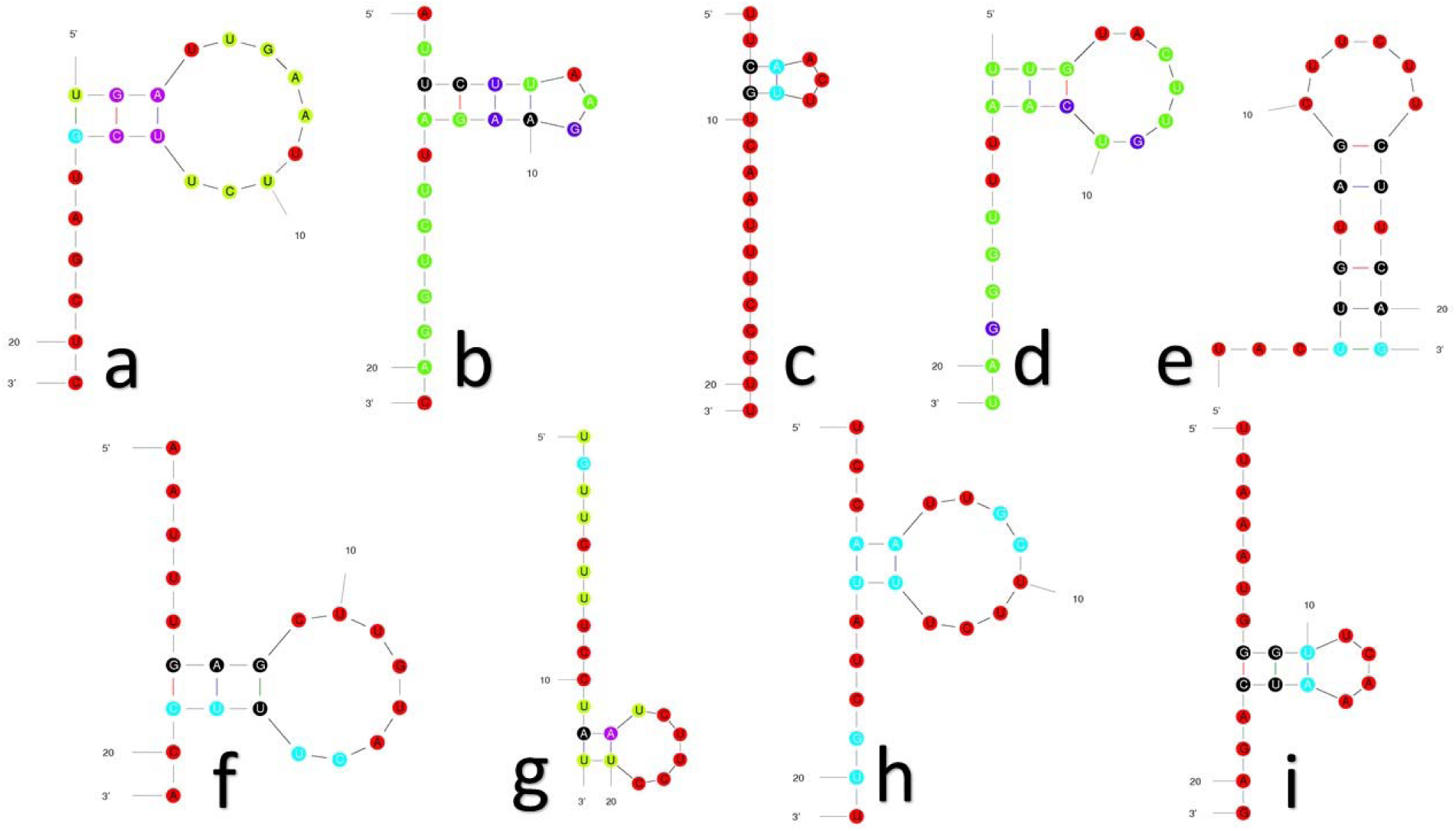
a-i, illustrating 9 different siRNA secondary structures from Mfold web server

**Figure 03:**
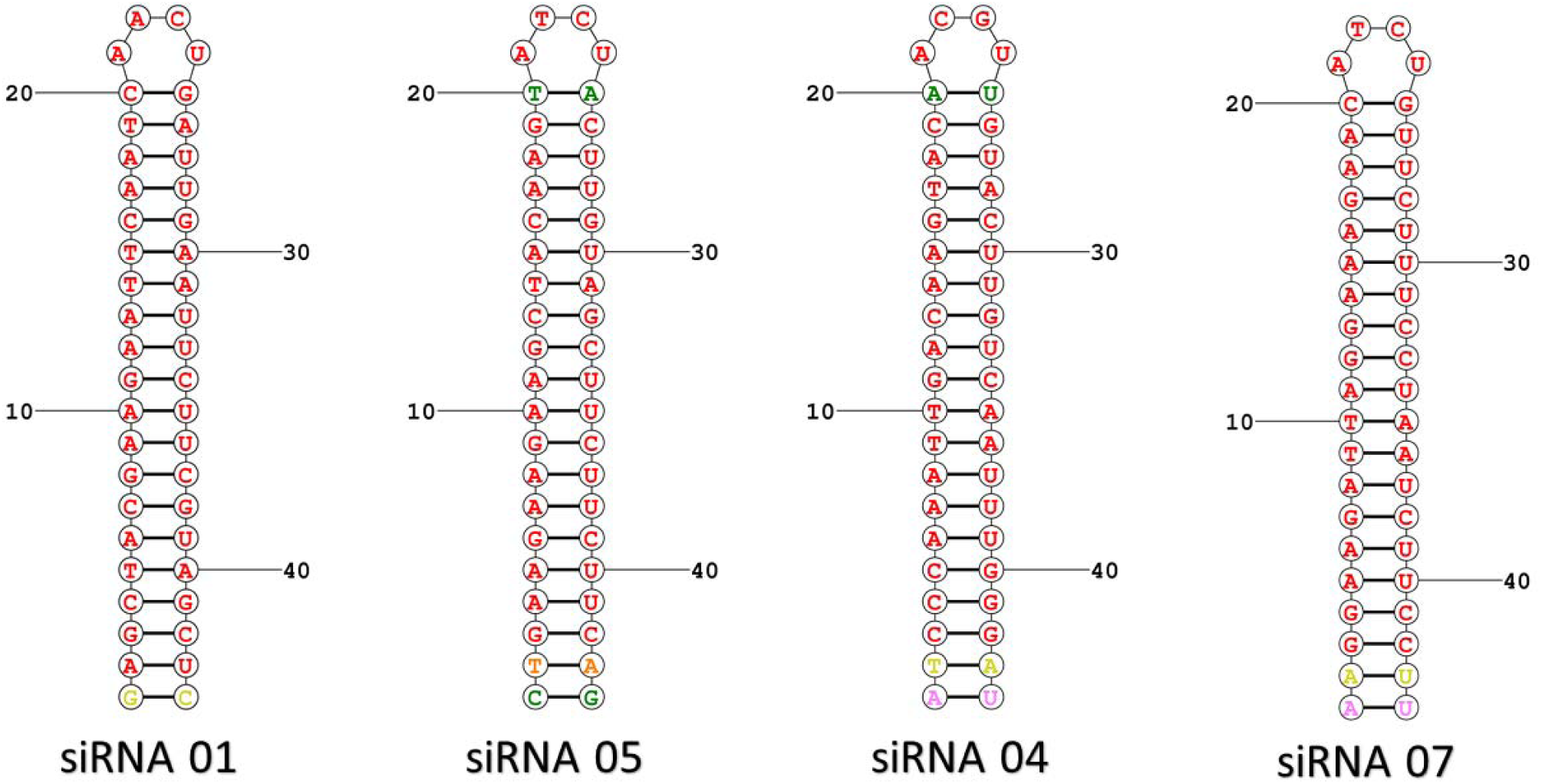
Predicted siRNA bonded with its corresponding target sequences forming a siRNA-target duplex secondary structure. Here, siRNA 01 and siRNA 05 possessed similar free energy of binding which is -31.4 kcal/mol where siRNA 04 and siRNA 07 possessed similar free energy of binding that is -30.0 kcal/mol.

**Figure 04:**
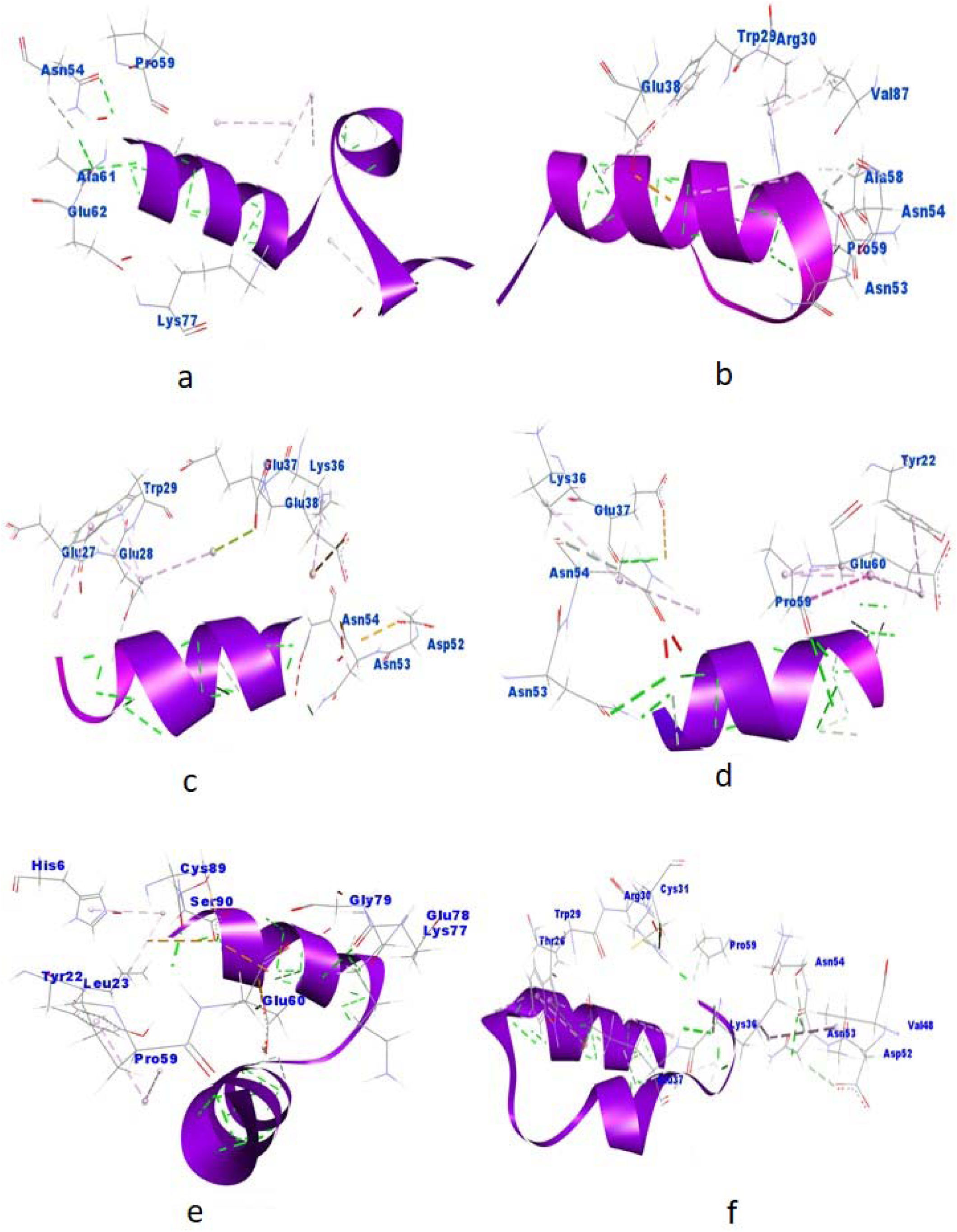
Molecular docking analysis of MSP-1 (19) with different anti-parasitic peptides (a to f). Here, (a), (c) and (e) depicts the docking complex of AP02283, AP00101 and AP00146 respectively with MSP-1. Moreover, Asn54, Lys36 and His6 were the only single amino acid of binding groove of MSP-1 where AP02283, AP00101 and AP00146 interacted respectively. Whereas (b) is showing the interaction of AP02285 with MSP-1 including major amino acids Arg30, Asn53, Asn54. Similarly, figure (d) and (f) are showing interactions with Lys36, Glu37, Asn53, Asn54 for both AP00190 and AP02284 respectively.

