## Supplemental File 1 for "An extensive computational approach to inhibit MSP-1 of *P.vivax* elucidates further horizon in the establishment next generation therapeutics against malaria"

Supplementary Table: Conserved Sequences from MSP-1 protein sequences of *P.vivax*

| 01 | ATGAAGGCGCTACTCTTTTTGTTCTCTTTCATTTTTTTCGTTACCAAATGTCAATGTGAAACAGAAAGTTATAAGCAGCTTGTAGCCAAGCTGGACAAGTTAGA |
| --- | --- |
| 02 | TGCAAATAATAATAATAACAATCAGGTTAGCGTTTTA |
| 03 | GAACCCAATGGGATAAAATACCTTGTGGAGAGCTACGAAGAATTCAATCAACTGATGCACGTGAT |
| 04 | TTGCTGGAAAAAATTATGAATAAAATTAAGATAGAAGAAGACAAATTGCCC |
| 05 | GACTTCTCCGGGGTTGTGGAATTACAAGTACAAAAGGTATTGATAATCAAAAAAATTGAGGCTCTAAAGAATGTCCAGAATCTTCTTAAGAATGCCAAGGTGAAGGACGACCTGTAC |
| 06 | GCGAGAAACCTGAGCCCTACTACTTGATGGTCCTCAAAAGGGAAATTGACAAGTTGAAGGA |
| 07 | CAGAACAACTTGCCAGCCATGTACTCCATATATGACTC |
| 08 | TGTACCAGAAGGAAATGGTTTACAATATATA |
| 09 | AAGCACCTATCCCAAATTGACAAGTACAACGA |
| 10 | AGAAATTGAGGAACTGAAGAAGAAGCTACAAGTATCTCTGGACCACTATGGAAAGTACAAGCTCAAATTGGAGAGG |
| 11 | TCCTCAAAAAGAAGAATAAAATCTCTAACAGCAAGGATCAAATTAAAAAGCTCACCAGTTTGAAAAACAAATTGGAGAGAAGACAAAATCTGTTGAATAACCCAACAAGTGTGTTGAAAAATTACACCGCTTTTTTCAACAA |
| 12 | AAGAGAGAAACAGAAAAGAAGGAGGTGGAAAATACCCTTAAGAATACCGAGATTTTGCTGAAGTACTATAAGGCACGAGCCAAATATTATAT |
| 13 | TGGAAGGAAGATTAGGAAAGAACATCGA |
| 14 | GCCTGAGATCCTCGTGCCAGCAGGAATCAGCGATTACGATGTGGTCTACTTAAAGCCATTAGCCGGAATGTACAAAACGATAAAGAAGCAATTGGAAAATCACGTAAACGCATTTAACACTAACATAACGGATATGTTAGACTCTAGACTGAAGAAGAGAAACTACTTCTTAGAAGTTCTGAACTCTGATTTGAACCCATTTAAGTATTCA |
| 15 | GAGAAAAAGGAAGCCGAGATCACTGTAAAGAAATTGCAGGACTACAACAAGATGGATGAGAAGTTGGAGGAGTACAAAAAATCGGAGAAAAAAAATGAAGTGAAGTCTTCTGGTCTTCTGGAAA |
| 16 | ATTGATGAAATCAAAATTGATTAAAGAAAACGAGTCCAAGGAAATATTATCCCAGCTGCTAAATGTGCAAACTCAGTTATTAACTATGAGCTCCGAGCACACATGTATAGACACCAATGTGCCTGATAATGCAGCCTGCTATAGGTACTTGGACGGAACGGAAGAATGGAGATGCTTGTTAACCTTTAAAGAAGAAGGCGGCAAGTGTGTGCCAGCATCGAATGTGACTTGTAA |
| 17 | AAATCGTCTGTAAATGTACTAAAGAAGGTTCTGAGCCACTCTTTGAGGGAGTTTTCTGTAGCTCCTCCAGCTTCCTAAGCTTGTCCTTCTTGTTGCTCATGTTGCTTTTCCTCCTGTGCATGGAGCTTTAA |
