## Supplemental File 2 for "An extensive computational approach to inhibit MSP-1 of *P.vivax* elucidates further horizon in the establishment next generation therapeutics against malaria"

| Sl | siRNA | U R A Rules | target position | contiguous G's or C's constraint | contiguous A's or T's constraint | GC content |
| --- | --- | --- | --- | --- | --- | --- |
| 01 | UGAUUGAAUUCUUCGUAGCUC; GCUACGAAGAAUUCAAUCAAC | URA | OK | OK | OK | OK |
| 02 | AUUCUUAAGAAGAUUCUGGAC; CCAGAAUCUUCUUAAGAAUGC | URA | OK | OK | OK | OK |
| 03 | UUCAACUUGUCAAUUUCCCUU; GGGAAAUUGACAAGUUGAAGG | URA | OK | OK | OK | OK |
| 04 | UCAUAUAUGGAGUACAUGGCU; CCAUGUACUCCAUAUAUGACU | URA | OK | OK | OK | OK |
| 05 | UUGUACUUGUCAAUUUGGGAU; CCCAAAUUGACAAGUACAACG | URA | OK | OK | OK | OK |
| 06 | UACUUGUAGCUUCUUCUUCAG; GAAGAAGAAGCUACAAGUAUC | URA | OK | OK | OK | OK |
| 07 | AAUUUGAGCUUGUACUUUCCA; GAAAGUACAAGCUCAAAUUGG | URA | OK | OK | OK | OK |
| 08 | UGUUCUUUCCUAAUCUUCCUU; GGAAGAUUAGGAAAGAACAUC | URA | OK | OK | OK | OK |
| 09 | UGUACAUUCCGGCUAAUGGCU; CCAUUAGCCGGAAUGUACAAA | URA | OK | OK | OK | OK |
| 10 | UCCAAUUGCUUCUUUAUCGUU; CGAUAAAGAAGCAAUUGGAAA | URA | OK | OK | OK | OK |
| 11 | UGUUAAAUGCGUUUACGUGAU; CACGUAAACGCAUUUAACACU | URA | OK | OK | OK | OK |
| 12 | UAAGAAGUAGUUUCUCUUCUU; GAAGAGAAACUACUUCUUAGA | URA | OK | OK | OK | OK |
| 13 | UUAAAUGGGUUCAAAUCAGAG; CUGAUUUGAACCCAUUUAAGU | URA | OK | OK | OK | OK |

Supplementary Table: siRNA’s that followed all the UAR rules and other rules.
